# Nest relief in the cryptically-incubating semipalmated sandpiper is quick, but vocal

**DOI:** 10.1101/2021.07.30.454457

**Authors:** Martin Bulla, Christina Muck, Daniela Tritscher, Bart Kempenaers

## Abstract

Biparental care requires coordination between parents. Such coordination might prove difficult if opportunities to communicate are scarce, which might have led to the evolution of elaborate and noisy nest relief rituals in species facing a low risk of predation. However, whether such conspicuous rituals also evolved in species that avoid predation by relying on crypsis remains unclear. Here, we used a continuous monitoring system to describe nest relief behavior during incubation in an Arctic-breeding shorebird with passive nest defense, the semipalmated sandpiper (*Calidris pusilla*). We then explored whether nest relief behavior provides information about parental cooperation and predicts incubation effort. We found that incubating parents vocalized twice as much before the arrival of their partner than during other times of incubation. In 75% of nest reliefs, the incubating parent left the nest only after its partner had returned and initiated the nest relief. In these cases, exchanges were quick (25s, median) and shortened over the incubation period by 0.1 – 1.4s per day (95%CI), suggesting that parents became more synchronized. However, nest reliefs were not cryptic. In 90% of exchanges, at least one parent vocalized, and in 20% of nest reliefs the incubating parent left the nest only after its returning partner called instantaneously. In 30% of cases, the returning parent initiated the nest relief with a call; in 39% of these cases, the incubating partner replied. If the partner replied, the next off-nest bout was 1 – 4hr (95%CI) longer than when the partner did not reply, which corresponds to an 8 – 45% increase. Our results indicate that incubating semipalmated sandpipers, which rely on crypsis to avoid nest predation, have quick but acoustically conspicuous nest reliefs. Our results also suggest that vocalizations during nest reliefs may be important for the division of parental duties.

**LAY SUMMARY:**

- Biparental care requires coordination between parents. In species where both parents incubate and nests are visible, parents often perform complex nest relief rituals. Is this also the case in species where nests are cryptic?
- We video-recorded nest relief behavior at nests of cryptically incubating semipalmated sandpipers.
- Parents relieved each other quickly, but they vocalized frequently. In 20% of nest reliefs the incubating parent left only after its returning partner called instantaneously. In 30% of cases the returning parent initiated the nest relief with a call. If the partner replied, its next off-nest bout was 1 – 4hr longer than when the partner did not reply, which corresponds to an 8 – 45% increase.
- Our results suggest that vocalizations during nest relief may be important for the division of parental duties. Further work in sandpipers and other taxa is needed to elucidate the role of parental vocalization in coordinating and synchronizing parental duties.

## INTRODUCTION

Biparental care requires cooperation and coordination between parents. Such need for coordination is high when only a single parent can care at a given time and when continuous care increases reproductive success. This situation occurs in birds during biparental incubation of eggs when parents coordinate their duties to ensure continuous nest attendance (Clutton-Brock 1991; Deeming 2002a).

Whereas instantaneous coordination of incubation duties is possible in species where the off-duty parent (i.e. the one taking an incubation recess) stays near the nest (Smith *et al.* 1978; Hawkins 1986; Glutz von Blotzheim 1999; Rodewald 2015; Sládeček *et al.* 2019), such coordination is challenging in species where the off-duty parent forages or rests out of calling distance of the nest (Bulla *et al.* 2015b). In some species, this can be hundreds or even thousands of kilometers away (Weimerskirch *et al.* 1993; Weimerskirch 1995; González-Solís *et al.* 2000; Boersma and Rebstock 2009). In such cases, communication among partners is often limited to a brief time when parents exchange on the nest.

Perhaps as a consequence of this limited possibility to communicate, some species – usually breeding in colonies and in environments with low risk of predation – developed elaborate nest relief rituals (Glutz von Blotzheim 1999; Deeming 2002b; Rodewald 2015). In contrast, the parents of species with cryptic nests and plumage that rely on passive nest defense (Moseley 1979; Glutz von Blotzheim 1999; Rodewald 2015) are expected to relieve each other on the nest quickly and discretely, because activity at the nest increases nest predation (Martin *et al.* 2000; Smith *et al.* 2012). However, little is known about nest relief behavior of such crypticaly incubating species and hence it is unclear how parents resolve the conflict between staying cryptic and communicating (coordinating) with their partner.

Here, we describe the nest relief behavior of a biparentally incubating shorebird, the semipalmated sandpiper (*Calidris pusilla*). This species breeds in the High Arctic and is cryptic during incubation, i.e. incubating parents often weave a ‘roof’ over the nest cup, sit tight when a predator flies by (Ashkenazie and Safriel 1979), and upon approach of a human often stay on the nest until nearly stepped upon (Bulla *et al.* 2016). A parent sits continuously on the nest for about 11 hours and then is quickly relieved by its off-duty partner (Bulla *et al.* 2014). Off-duty parents forage up to 2 km from their nest; thus, instantaneous vocal communication between the incubating parent and its off-duty partner is unlikely (Bulla *et al.* 2015b).

Using video recordings and an automated incubation monitoring system (Bulla et al. 2014), we investigated whether the incubating parent actively initiates the nest relief by calling its partner to the nest or by flying off the nest to inform its partner of its need to be relieved. Next, we quantitatively and qualitatively described the actual nest relief process. As semipalmated sandpipers were surprisingly vocal during nest relief, we further explored whether vocalizations during nest relief provide information about parental coordination. Specifically, we tested (1) whether the duration of nest relief and the intensity of vocalizations during nest relief decrease as parents become more synchronized over the incubation period (Bulla *et al.* 2014), and (2) whether the intensity of vocalizations conveys information, i.e. correlates with the length of the current incubation bout or – similar to zebra finches, *Taeniopygia guttata* (Boucaud *et al.* 2016a) – predicts the length of the next incubation bout.

## METHODS

### Data Collection

We monitored the incubation behavior of semipalmated sandpipers near Utqiagvik, Alaska (71.32°N, 156.65°W) during June and July in 2011 to 2013. The breeding area and the procedures of nest searching, catching of parents, as well as the monitoring of incubation have been described in detail elsewhere (Ashkenazie and Safriel 1979; Bulla *et al.* 2014; Bulla *et al.* 2015a).

In short, we used a custom-designed video recording system consisting of an external optical lens (ø 2 cm, length 4 cm) mounted on a tripod and placed 1 - 3 meters from the nest (Appendix Picture A1) and an external microphone (ø 0.75 cm, length 2.2 cm) placed in the vegetation about 1 meter from the nest. Both were connected to a recorder hidden in the vegetation about 5 meters from the nest. The system was powered by a 12-V, 31-Ah or 44-Ah battery hidden in the vegetation, and allowed continuous recording for 1 - 3.5 days. Each parent was banded with a unique combination of color bands and with a green plastic flag that contained a passive tag (9.0 mm × 2.1 mm, 0.087g, Biomark, Boise, Idaho 83702, USA, http://www.biomark.com/). The tag, and hence the presence of a parent on the nest, was registered throughout the incubation period every 5 s by a radio frequency identification device with a thin antenna loop around the nest cup and connected to a reader. We used these data (a) to verify the identity of the parents, (b) to identify the time of nest relief, and (c) to extract the length of each incubation bout - the time between the incubation start of a parent and its departure from the nest followed by incubation of its partner (Bulla *et al.* 2014; Bulla *et al.* 2015a). Note that the incubation bout of a parent represents the off-duty bout of its partner. At 3 nests (11 nest relief observations) the radio frequency identification system temporarily failed, but nest reliefs were identified directly from the video recordings using the color bands of the parents.

Some nests were protected from avian predators, at least for some time, by cages made of mesh wire (*N*_2011_ = 0 out of 5 nests, *N*_2012_ = 12 out of 14, and *N*_2013_ = 8 out of 12 nests; for cage designs see Appendix Picture A1 and figure S2 in Bulla et al. 2019). The cages may have affected the nest relief behavior. For example, the incubating parent could not directly fly off the nest, but had to walk out of the cage first.

### Extraction of Nest Relief Behavior from Video-recordings

From the video recordings, we noted all cases of an incubating bird being relieved by its partner (*n* = 163). Specifically, we noted when the incubating parent left the nest (*departure*), which occurred either prior to or after the return of its partner to the vicinity of the nest (*arrival*). We also noted how it left the nest (flew off, walked off, walked off and then flew off). To detect the *arrival* of the returning parent, we screened video recordings starting 30 min before the incubating parent left the nest. We also noted when the returning parent started walking towards the nest (*initiation*) and when it sat down on the nest (*incubation start*).

To investigate whether the incubating parent changed its behavior shortly before its partner arrived, we compared the calling and the fly-offs from the nest (defined in Table 1) during the period starting 30 min before the incubating parent left the nest and ending with the *arrival* of its partner near the nest (*before-arrival* observation) and during a control period randomly selected from the recordings, such that it started the earliest 35 min after incubation started and ended at least 60 min prior to the end of the incubation bout of the focal parent (*control* observation). For each *before-arrival* and *control* observation, we noted whenever the incubating parent called or flew off the nest (Table 1). In total, we quantified behavior from 125 *before-arrival* observations (i.e. observations where the incubating parent left only after the returning parent *arrived* and calling could be scored; median = 3, range: 1-14 observations per nest) and from 114 *control* observations (median = 5, range: 1- 5 observations per nest) from 29 nests.

**Table 1.**
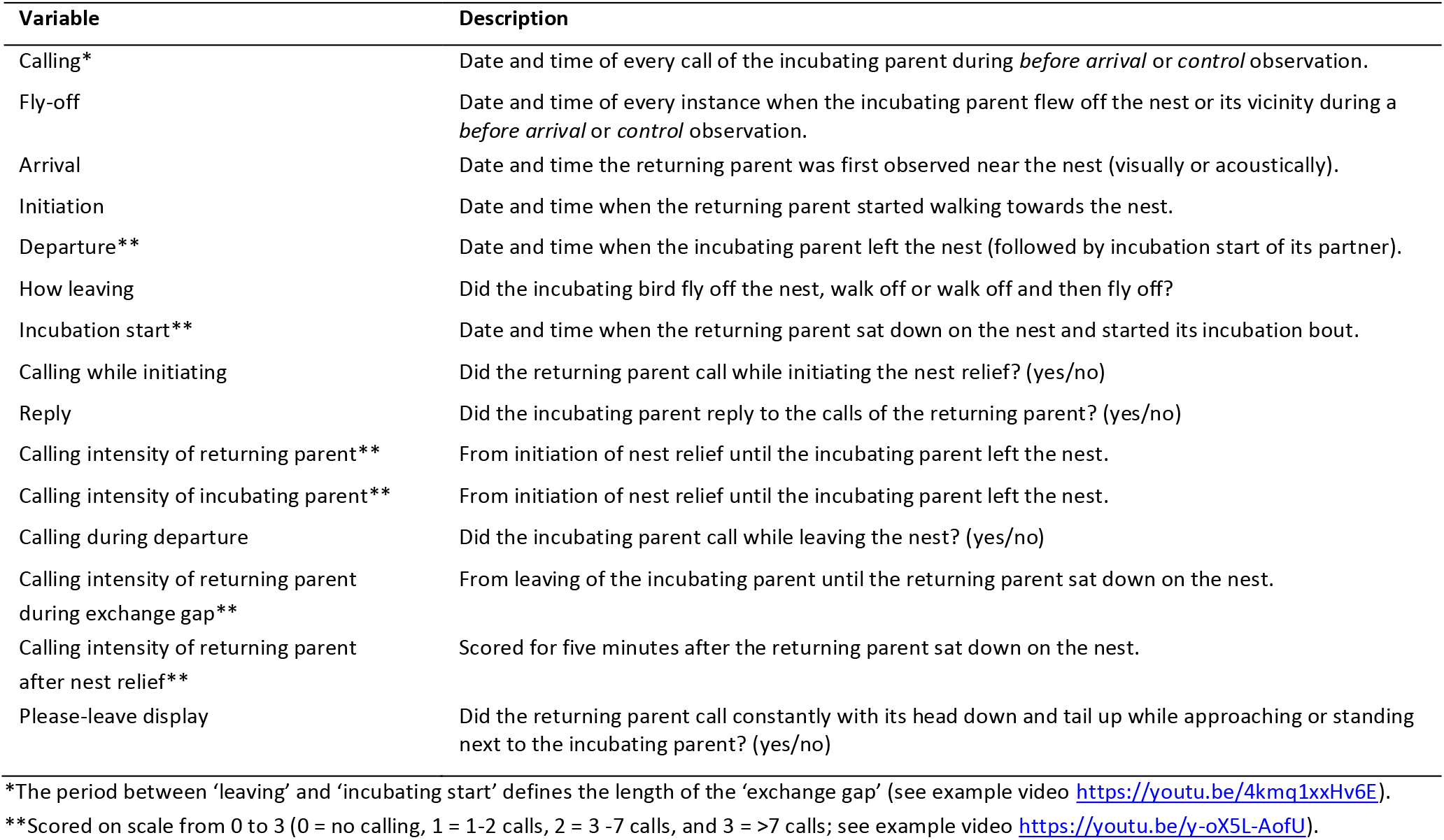
Nest relief parameters extracted from the video recordings.

To investigate the timing of actual nest relief, we noted (1) whether the incubating parent left before the returning parent arrived (yes/no), as such nest reliefs apparently do not involve communication between the partners, and (2) for how long the eggs were left unattended (*exchange gap*), i.e. the period between the incubating parent’s *departure* and the *incubation start* of the returning parent. For nest reliefs where the incubating parent left only after the returning parent arrived, we further noted (3) the latency of the returning parent to initiate the nest relief (*latency to initiate*), i.e. the period between its *arrival* and nest relief *initiation*, and (4) the length of the period from nest relief *initiation* until *departure* of the incubating parent (*from initiation to departure*).

We further investigated vocalizations during and after nest relief. From the video recordings, we noted (1) whether the returning parent called when initiating the nest relief (yes/no), (2) whether its incubating partner replied (yes/no), and (3) whether the incubating parent called during *departure* from the nest (yes/no). We also scored calling intensity on a scale from 0 to 3 (Table 1) (4) of both parents during the period from *initiation* to *departure*, (5) of the returning parent during the *exchange gap*, and (6) during the 5 min after it started incubating. In addition, we noted (7) every instance where the returning parent called constantly with its tail up while approaching or standing next to the incubating parent (’*please-leave’ display*; yes/no; see video https://youtu.be/iIBwFQPreDw).

All behavioral parameters are summarized in Table 1. Examples of video recordings illustrating each behavior can be found at https://bit.ly/38vBOhC. Videos were scored by C.M. and D.T., blindly to the sex of the birds. For some observations not all behaviors could be scored (e.g., calling, because of poor sound quality); thus, sample sizes vary between statistical tests.

### Statistical Analyses

We used R version 3.5.2 (R-Core-Team 2020) for all statistical analyses and the *lme4* package (Bates *et al.* 2015) for fitting the mixed-effect models. We used the *sim* function from the *arm* package and a non-informative prior distribution (Gelman and Hill 2007; Gelman and Su 2018) to create a sample of 5,000 simulated values for each model parameter (i.e. posterior distribution). We report effect sizes and model predictions by the medians, and the uncertainty of the estimates and predictions by the Bayesian 95% credible intervals represented by the 2.5 and 97.5 percentiles (95%CI) from the posterior distribution of 5,000 simulated or predicted values. We graphically inspected the goodness of fit, and the distribution of the residuals. All results are reproducible with the open access data and code available from the Open Science Framework (Bulla 2019), which also provides visual representations of model assumptions.

## RESULTS

### Behavior of the Incubating Parent Before the Return of Its Partner

*Before arrival* of the off-duty parent, the calling rate of the incubating parent was twice as high than during other times during its incubation (*before arrival*: 0.4, 95%CI: 0.3 – 0.6 calls per 10 min; *control*: 0.2, 0.1 – 0.3; Figure 1a, Appendix Table A1a, b). In contrast, the probability that the incubating parent flew off the nest was similar (*before arrival*: 0.11, 0.07 – 0.19; *control*: 0.10, 0.05 – 0.17; Figure 1b, Appendix Table A1c, d). The calling rate and the probability of fly-offs did not differ between the sexes, and did not change over the incubation period (Appendix Table A1b, d).

**Figure 1.**
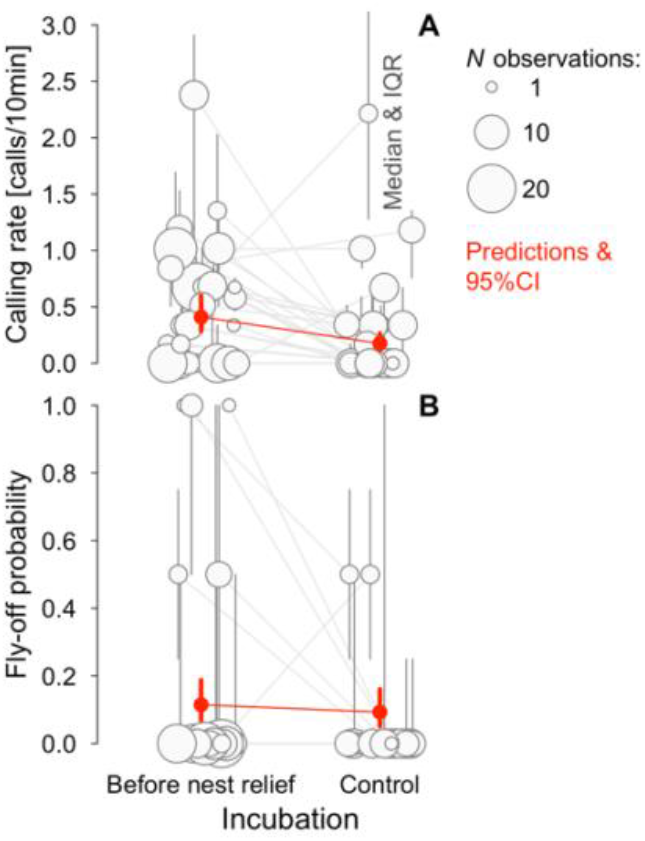
Calling rate (A) and the probability of flying off the nest by the incubating parent (B) *before arrival* of its off-duty partner and during other times of incubation (*control*). Circles with bars indicate median and interquartile range for each of 29 nests. Circle size indicates the number of nest reliefs. Grey lines connect observations from the same nest. Red points with bars indicate model predictions with 95%CI based on the joint posterior distribution of 5,000 simulated values generated from the model output (Appendix Tables A1a, c) by the *sim* function in R (Gelman and Su 2018), while keeping the other predictors constant. *n* = 125 *before-arrival* and 114 *control* observations.

### Nest Relief Behavior

#### Timing

In general, the off-duty parent arrived in the vicinity of the nest shortly before it initiated the nest relief (median = 0 s, mean = 26 s, range: 0 - 19 min before nest relief initiation, *n* = 163 nest reliefs at 31 nests); this did not differ between the sexes and did not change over the incubation period (Appendix Table A2a). In 76% of nest reliefs the incubating parent left the nest by directly flying off (47 cases out of 62 nest reliefs without cage), in 18% by walking off and then flying (11 cases), and in the remaining 6% by walking off and out of sight of the camera (4 cases). In 15% of nest reliefs the incubating parent had left the nest before the *arrival* of the off-duty parent (25 cases from 15 nests out of 163 nest reliefs from 31 nests). However, in only 10 of these 25 cases did the incubating parent leave the nest >1 min prior to its partner’s return (median = 4.6 min, mean = 40 min, range: 2.6 min – 3.9 h, *n* = 10; in 3 cases >10 min), which suggests that in the remaining 15 cases the returning partner might have been already around, but had not been detected based on the video recording. In a further 9% of nest reliefs (15 cases from 11 nests), the incubating parent left the nest while the returning parent was around, but had not yet initiated the nest relief. Accordingly, in 75% of nest reliefs (123 cases from 30 nests) the incubating parent left only after its partner initiated the nest relief.

Once the returning parent initiated the nest relief, it took on average only 22 s until it started incubating (median, mean = 25 s, range: 4 s – 1.9 min, *n* = 163 nest reliefs at 31 nests); this did not differ between the sexes and across the incubation period (Appendix Table A2b). In the 123 cases where the incubating parent left only after its partner initiated the nest relief, the returning parent took on average 15 s (95%CI: 14 – 16 s) longer to start incubating than in the 40 cases (25 + 15) where the incubating parent left before the returning partner initiated the nest relief (Figure 2, Table A2b). Moreover, in 20% of the 123 nest reliefs (i.e. in 25 cases at 13 out of 30 nests) the incubating parent only left the nest after the returning parent called incessantly while approaching or standing right next to the nest with its head down and tail up, in an apparent attempt to convince its mate to leave (*please-leave display*). In all of the 13 nests where the *please-leave display* occurred, either only the female (8 nests) or only the male (5 nests) performed the display. In other words, we found no nest where both parents performed the display. The *please-leave display* was more common when the male was incubating (28% - 18 out of 65 cases) than when the female was incubating (12% - 7 out of 58 cases), but the difference was not significant when controlling for bird identity (Table A3). The period between the *initiation* of nest relief by the returning parent and the *departure* of the incubating partner shortened over the incubation period by about 0.1 – 1.4 s per day (95%CI; Figure 3a, Table A2d) and was 2.5 times (18 s) longer in nest reliefs with *please-leave display* than in those without such a display (95%CI: 12 – 24 s; Figure 3b, Table A2d). The probability of *a please-leave display* was independent of the length of the current incubation bout, but increased somewhat over the incubation period (Table A3).

**Figure 2.**
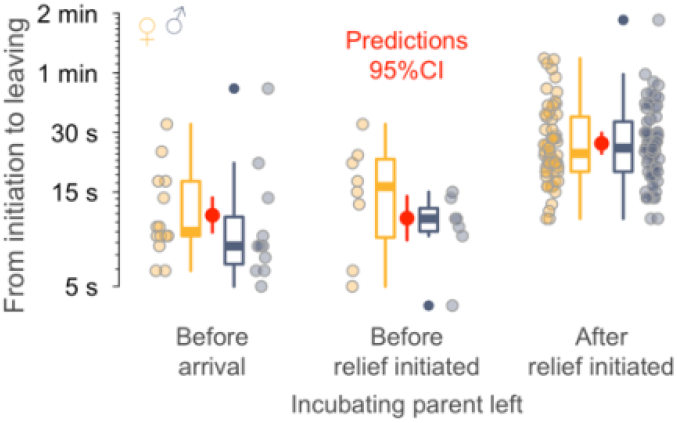
Period from nest relief *initiation* to *incubation start* according to when the incubating parent left the nest. The incubating parent left before the returning partner arrived (before arrival), while the returning partner was around, but had not yet initiated the nest relief (while around) or after the returning partner initiated the nest relief (after relief initiated). Circles indicate single observations. Box plots depict median (horizontal line inside the box), the 25^th^ and 75^th^ percentiles (box), the 25^th^ and 75^th^ percentiles ±1.5 times the interquartile range or the minimum/maximum value, whichever is smaller (bars), and the outliers (dots). Color of circles and box plots indicates the sex of the returning parent (female = yellow, male = blue-grey). Red points with bars indicate model predictions with 95%CI based on the joint posterior distribution of 5,000 simulated values generated from the model output (Appendix Table A2b) by the *sim* function in R (Gelman and Su 2018), while keeping the other predictors constant. *n* = 163 observations from 31 nests.

**Figure 3.**
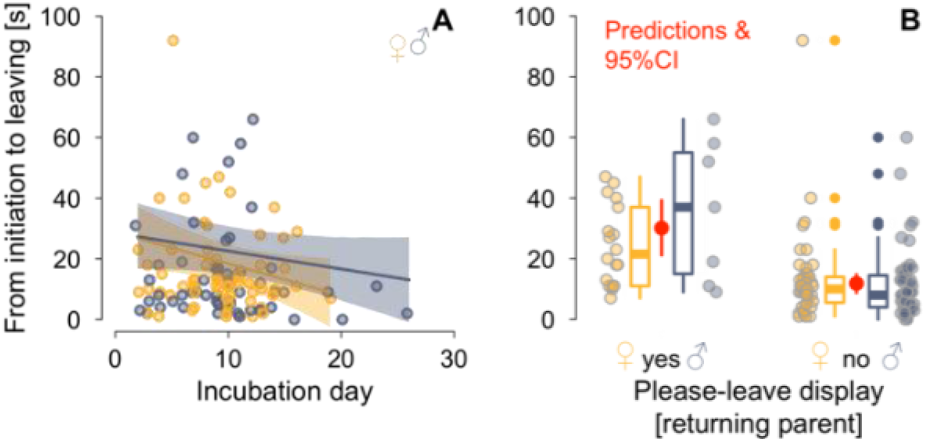
Period between nest relief *initiation* until *departure* in relation to the progress of the incubation period (A) and in relation to the occurrence of the *please-leave display* (B). Circles indicate single observations. Color indicates the sex of the returning parent (female = yellow, male = blue-grey). Lines with shaded areas (**A**) and red points with bars (**B**) indicate model predictions with 95%CI based on the joint posterior distribution of 5,000 simulated values generated from the model output (Appendix Table A2d) by the *sim* function in R (Gelman and Su 2018), while keeping the other predictors constant. Box plots are defined in the legend of Figure 2. *n* = 123 observations (from 30 nests) of an incubating parent leaving only after the returning parent initiated the nest relief.

The exchange gap, the time between *departure* of the incubating parent and *incubation start* of the returning partner, took on average 15 s (median, mean = 2.8 min, range: 0 s - 4 hr, *n* = 163 nest reliefs at 31 nests) and this was not sex specific and did not change over the incubation period (Table A2c). The exchange gaps were longest when the incubating parent left before its partner arrived (median = 1.4 min, mean = 17 min, range: 13 s – 4 hr, *n* = 25 nest reliefs at 14 nests) and shortest in the 123 cases where the incubating parent left only after its partner initiated the nest relief (median = 13 s, mean = 14 s, range: 0 – 36 s; Table A2c).

#### Vocalizations

While initiating the nest relief the returning parent called in 30% of all observations (46 out of 155 cases where calling could be scored, 32 out of 117 cases when the incubating parent left only after the returning parent initiated the nest relief). In 39% of those cases the incubating parent replied to the call of its returning partner (12 out of 31 cases where reply could be scored; 5 males from 5 nests and 7 females from 7 nests). The probability to call or reply was similar for both sexes and did not change over the incubation period (Appendix Table A4a, b).

Although calling varied widely between individuals and between nest-relief events (Appendix Figure A1, Table A4), in general, the incubating parent was less vocal during nest relief than the returning parent (Figure 4a). The returning parent called more often during the period when its incubating partner was still on the nest (Figure 4a & A1), less often after its partner had left (i.e. during the exchange gap, Figure 4b & A1), and was mostly quiet after sitting down on the nest (Figure 4c & A1). If the returning parent vocalized, it did so throughout most of the nest-relief process (Figure A1), but in 20% of nest reliefs the returning parent was completely silent (26 out of 129 cases), even when its partner was on the nest upon *arrival* (14%, 15 out of 108 cases; Figure A1). The calling intensity of the incubating and returning parent correlated positively between *initiation* and *departure* (Table A4c). The calling intensity of the returning parent was independent of sex and did not change over the incubation period (Table A4). The incubating parent called while leaving the nest in 19% of nest reliefs (28 out of 150 cases where calling could be scored; in 25% of nest reliefs without cage - 15 out of 60) and males were twice as likely to call (25% - 19 out of 75 cases, 34% - 11 out of 32 of cases without cage) than females (12% - 9 out of 75 cases, 14% - 4 out of 28 of cases without cage); the probability that the incubating parent called when departing did not change over the incubation period (Table A5). The incubating parent was silent in 64% of nest reliefs (54 out of 83 cases where the incubating parent left after *initiation* and all its vocalizations could be scored). In 10% of nest reliefs neither parent called (8 out of 83 cases where incubating parent left after *initiation* and all vocalizations could be scored).

**Figure 4.**
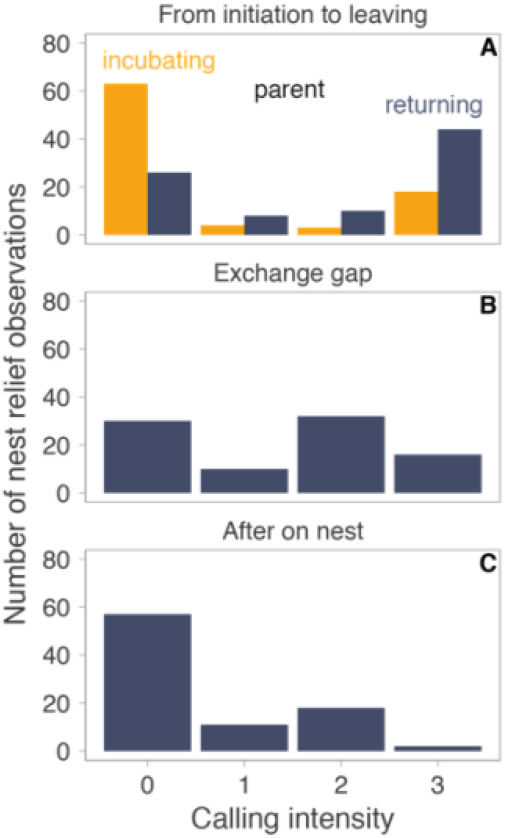
Distribution of calling intensities according to the stage of the nest relief. **A,** Calling intensity of incubating (yellow) and returning parent (blue) from the *initiation* of nest relief by the returning parent until the incubating parent left the nest, i.e. when both parents were present. **B,** Calling intensity of the returning parent during the exchange gap, i.e. after the incubating parent left the nest, but before the returning parent sat on the nest. **C,** Calling intensity of the returning parent during the 5 min after it started incubation. *N* = 88 nest reliefs (where all calling intensities were scored and where the incubating parent left only after the *arrival* of its partner) from 24 nests.

Calling intensities were unrelated to the length of the current and of the subsequent incubation or off-nest bout (Appendix Table A6 & A7). However, if the incubating parent replied to the call of its returning partner, the next off-nest bout was 1 – 4hr (95%CI) longer than when the incubating parent did not reply, and this effect was stronger when the incubating bird was the female (Figure 5a, Table A7b). A female reply to the returning male’s call was associated with a 42% (95%CI: 16 – 75%) increase in the length of the next incubation bout, i.e. an increase of 4hr (1.6 – 6.8hr). Furthermore, an increased calling intensity of returning females (but not males) during the 5 minutes after they sat down on the nest to incubate tended to be associated with a longer next incubation bout, albeit these estimates were noisy (Figure 5b; Table A7f).

**Figure 5.**
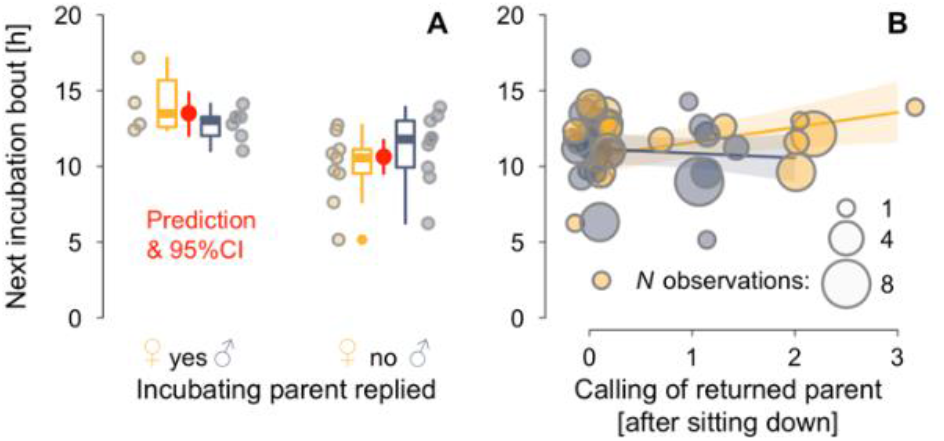
Calling behavior and the length of the incubation bout following the nest relief. The length of the next incubation bout depends on whether the incubating parent replied to the call of the returning partner emitted while initiating nest relief (**A**) and on the calling intensity of the returned parent during the 5 min period after it sat down on the nest to incubate (**B**). Circles in **A** indicate single observations, in **B** averages per individual with circle size indicating number of observations. Color indicates sex for x-axis variable (female = yellow, male = blue-grey). Red points with bars (**A**) and lines with shaded polygons (**B**) indicate model predictions with 95%CI based on the joint posterior distribution of 5,000 simulated values generated from the model outputs (Appendix Tables A7b, f) by the *sim* function in R (Gelman and Su 2018). *N*_a_ = 28 observations (of 22 parents from 17 nests) where the returning parent called before nest relief. *N***_b_** = 104 observations (of 44 parents from 26 nests) where the incubating parent left only after its partner initiated the nest relief. Definition of box plots is in Figure 2.

## DISCUSSION

We found that the general nest relief procedure of semipalmated sandpipers was brief, but vocal. When the incubating parent left only after its returning partner initiated the nest relief, the nest relief period shortened over the incubation period, suggesting that parents became more synchronized. We found some evidence that vocal communication between the incubating and returning parent contains information about parental investment.

### Nest Relief Procedure

The incubating parent called more often in the period before its partner arrived than at other times during incubation (Figure 1). Note, however, that we determined the *arrival* of the off-duty parent acoustically or visually from the video recordings, which have a limited field of view. We therefore cannot exclude that the off-duty parent had actually arrived near the nest earlier, which could explain the increased calling rate in the *before-arrival* observations.

Generally, but not always, the incubating parent waited for its partner to return before leaving the nest (85% of cases), mostly by flying directly off the nest. The returning parent was usually present in the nest vicinity just before the actual nest relief, and the nest relief procedure was brief (25 s on average) and vocal (Figure 4 and Appendix Figure A1). The return of the off-duty parent shortly before the nest relief reflects the previous finding that the incubating parent seems to wait for its partner to return before leaving the nest (Bulla *et al.* 2015b). The alternative explanation, that the incubating parent can accurately predict its partner’s return, seems less likely. Indeed, the exchange gaps were longer (up to 4h) when the incubating parent left before its partner arrived. The briefness of the actual nest relief also fits with previous findings that exchange gaps in semipalmated sandpipers, as well in other shorebirds, are short (Bulla *et al.* 2014; Bulla *et al.* 2016). Our results also corroborate the findings from shorebirds and seabirds with long incubation bouts (>12 h), showing that the incubating parent typically waits for the return of its partner until its energy stores become depleted (Davis 1982; Ball and Silver 1983; Chaurand and Weimerskirch 1994; Weimerskirch 1995; Gauthier-Clerc *et al.* 2001; Bulla *et al.* 2015a; Bulla *et al.* 2015b; Bulla *et al.* 2017). Thus, in these birds, the off-duty parent plays a decisive role in determining the length of an incubation bout.

### Vocalization and Cooperation

Nest relief events were surprisingly vocal (Figure 4 and Appendix Figure A1; for example, see video at https://youtu.be/y-oX5L-AofU). At least one parent vocalized in 90% of cases where the incubating parent left after its partner had initiated the nest relief and all calling parameters could be scored (*n* = 83). In general, the returning parent initiated the nest relief silently (70% of cases), but then vocalized until its partner left the nest (70% of cases), and vocalized also during the exchange gap, albeit with lower intensity (66% of cases, Figure 4). The returning parent then remained silent for 5 min after starting incubation (65% of cases, Figure 4c). The incubating parent was mostly silent during the nest relief (72% of cases) and left with a call in only 19% of cases. In 20% of nest relief events the incubating parent only left the nest after the returning parent started calling incessantly while approaching or standing right next to the nest (for video see https://youtu.be/iIBwFQPreDw), which prolonged the nest relief (Figure 3) and suggests that the incubating parent was reluctant to leave.

Such intense vocalizations during nest relief are surprising given the cryptic incubation of semipalmated sandpipers. However, because semipalmated sandpiper parents usually relieve each other only twice a day (Bulla *et al.* 2014) and because the off-duty parent typically forages away from the nest area (Bulla *et al.* 2015b), the nest relief period is likely the only occasion during which parents can communicate. Thus, the intense vocalizations during nest relief may play a role in this communication, and may be similar to the more elaborate nest relief rituals observed in other avian species (Glutz von Blotzheim 1999; Wachtmeister 2001; Deeming 2002b; Rodewald 2015). Vocalizations during nest relief might be crucial for synchronizing parental duties (Wachtmeister 2001; Servedio *et al.* 2019), but whether they indeed facilitate coordination among parents in semipalmated sandpipers remains to be shown. On the one hand, the vocalizations during nest relief events were not indicative of the length of a given incubation bout (Appendix Table A3 & A6), the intensity of the vocalizations did not decrease over the incubation period (Table A4 & A5), and the probability of the *please-leave display* may even have increased (Table A3). On the other hand, the interval between the *initiation* of nest relief and the *departure* of the incubating parent shortened over the incubation period (Table A2d), suggesting that parents became more synchronized over time and minimized the time they both spent at the nest. We also found two pieces of evidence that the calls may convey information about the length of the next incubation bout (Figure 5, Table A7). (1) When the returning parent initiated the nest relief with a call (30% of cases), and the incubating parent replied (39% of those cases), the next incubation bout of the returning parent, which is also the next off-nest bout of the previously incubating parent, was 1 - 4 hr longer than when no such reply occurred (Figure 5a). We thus hypothesize that the reply to a returning parent’s call signals the relieved parent’s need to stay longer off the nest than usual. This hypothesis can be tested, because it predicts (i) that the reply should be associated with a current poorer condition of the relieved parent (e.g., a lower body mass), and (ii) that experimental manipulation of the body mass (e.g., via supplemental feeding) should decrease the probability of reply. (2) The calling intensity of the returning parent during the 5-min period after it started incubating positively correlated with the next incubation bout. However, this was only the case in females, not in males (Figure 5b). Whether this effect is biologically relevant (and not a false positive) requires further study (replication).

Only few studies have linked vocalizations between the partners during incubation to parental investment. Pairs of lesser black-backed gulls (*Larus fuscus*) that vocalized for longer during nest relief seemed to divide the incubation duties more equally (Kavelaars *et al.* 2019). In northern lapwings (*Vanellus vanellus*), exchange gaps were shorter when males relieved their calling female than when they relieved a non-calling female, and whenever a calling female left the nest for an incubation recess (i.e. no nest relief took place), her incubation recess was longer than when she left for the recess silently (Sládeček *et al.* 2019). In zebra finches (*Taeniopygia guttata*), the calling rate of a returning male predicted the subsequent off-nest time of its female partner (Boucaud *et al.* 2016a), while calling of an incubating female from inside the nest predicted whether or not the male relieved her from the nest (Boucaud *et al.* 2017). Perhaps such vocalizations honestly signal the incubating bird’s needs, as suggested by both observations and experiments in some uniparentally incubating females, which show that females signal the need to be fed to their mate (Boucaud *et al.* 2016b; Boucaud *et al.* 2016c; reviewed in Amy *et al.* 2018).

### Conclusions

The brief, but vocal, nest relief events in a cryptically incubating shorebird seem puzzling, because vocalizations provide a clear indication of a nest. Yet, in species where brief nest reliefs are the only possibility for parents to communicate, selection might still favor vocalizations if those lead to better coordination between the parents and if the risk of detection by a predator remains low. Indeed, we found some evidence that vocalizations are associated with the immediate (next) investment of the partner. Given the lack of knowledge about vocalizations during biparental care, further work in this and other taxa is needed to elucidate the role of parental vocalization in coordinating and synchronizing parental duties.

## ACKNOWLEDGEMENTS

We thank A. Rutten, E. Stich, F. Heim, F. Prüter, I. Steenbergen, K. Chmel, K. Murböck, L. Langlois, L. Verlinden, M. Schneider, M. Šálek, M. Valcu, S. Herber, and V. Kubelka for help in the field, J. Petru for the custom-designed video-recording system, A. Girg for genetic sexing, and M. Valcu for advice on data analysis. This work was part of the PhD project of M.B. in the International Max Planck Research School for Organismal Biology.

## Ethics statement

The U.S. Department of the Interior, U.S. Fish and Wildlife Service, and State of Alaska Department of Fish and Game permitted the data collection (permit no. MB210464-0, MB210464-1, 2350, 23520, 11-106).

## Authors’ contributions

M.B. and B.K. conceived the study; M.B. with help of B.K. collected the data. C.M. with help of D.T. and M.B. extracted the behavior from video recordings. D.T., M.B., and C.M. extracted the length of incubation bouts. D.T. with help of M.B. prepared the data for analyses. D.T. with help of C.M. prepared the video examples. M.B. analyzed the data, prepared the supporting information and with help of B.K., C.M. and D.T. wrote the paper. All authors contributed to finalizing the paper.

## Data availability

All results reported in this article are reproducible with the open access data and code available from the Open Science Framework https://doi.org/10.17605/OSF.IO/E2W7Y (Bulla 2019), which also contains video examples and plots of model assumptions.

## APPENDIX

**Picture A1.**
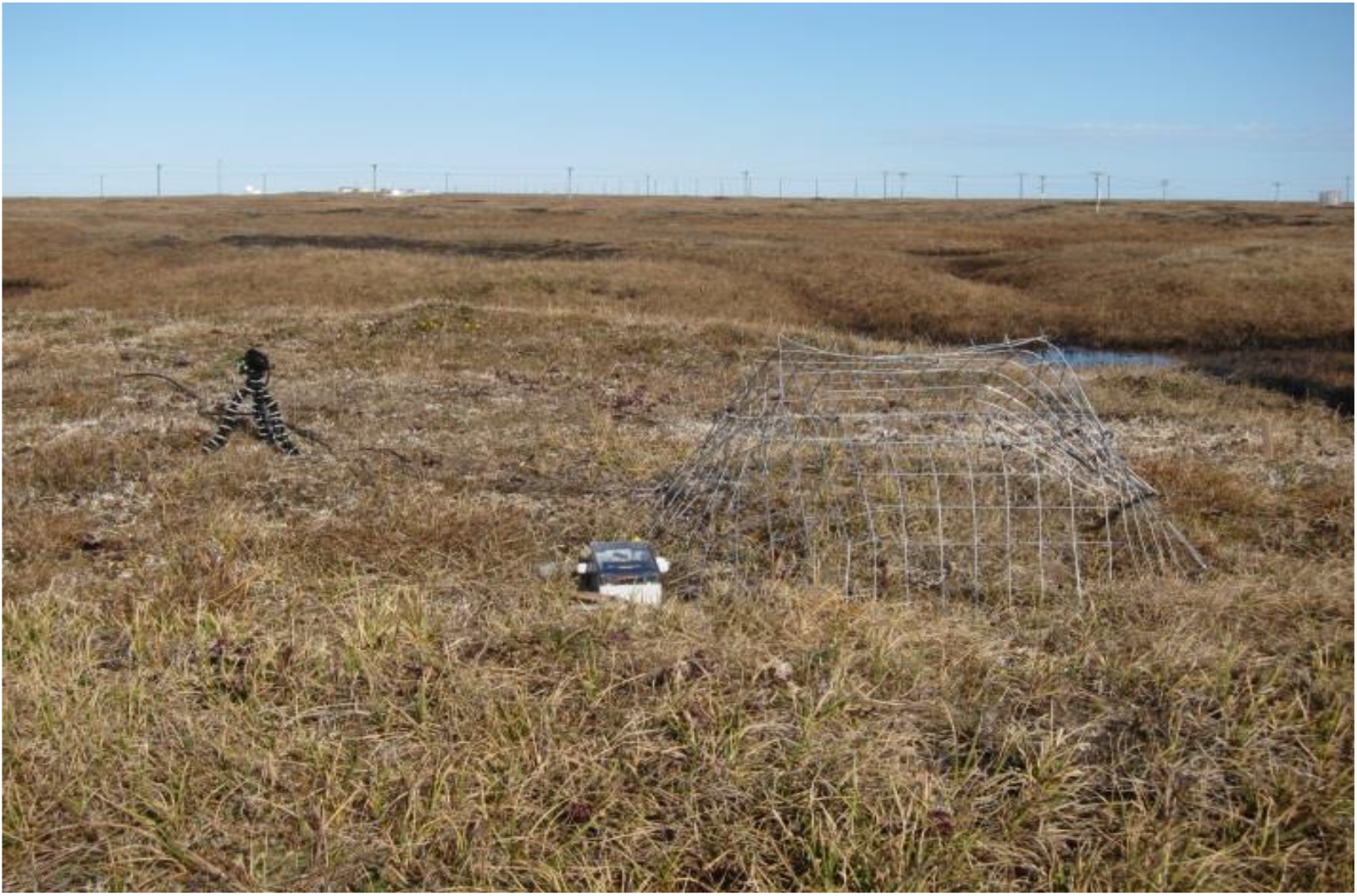
Recording equipment at the nest. The camera is mounted on a little tripod. The battery is hidden at the vegetation. Here, the nest is protected by a cage and radio frequency identification system is visible next to the cage.

**Figure A2.**
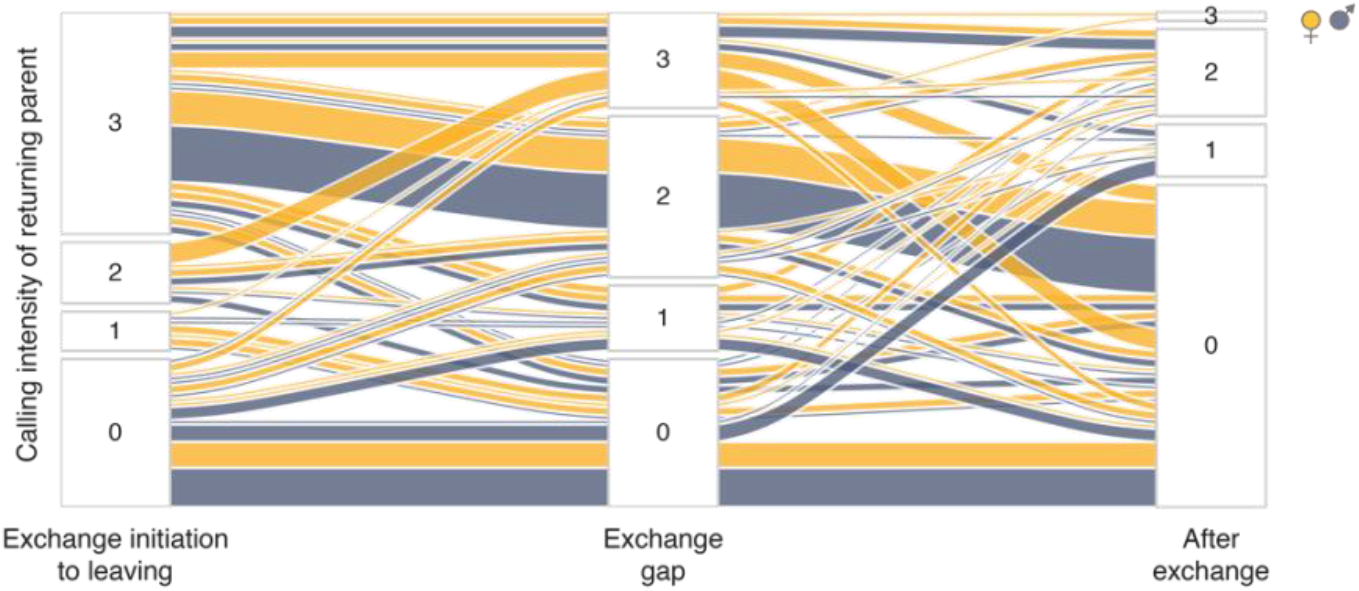
Calling intensities of the returning parent throughout the nest relief process. The numbers in the stacks indicate calling intensities, the sizes of the stacks indicate distribution of calling intensities of the returning parent across three time periods: from the *initiation* of nest relief until *leaving* of the incubating parent, i.e. when both parents were present (**left**), during the exchange gap, i.e. after the incubating parent left, but before the returning parent sat on the nest (**middle**), and during the 5 min after the returning parent started incubating (**right**). Each line represents different calling scenario, line colour the sex of the returning parent (female in yellow, male in blue-grey), line thickness indicates prevalence of the given scenario. Created using *alluvial* function from *alluvial* R package (Bojanowski and Edwards 2016). *N* = 108 nest reliefs where calling intensities could have been scored for all three periods and where the incubating parent left only after the *arrival* of its partner. Note that calling intensities of the returning parent throughout the nest relief process greatly varied, but the two most common scenarios were intense vocalization or being quiet throughout the nest relief.

**Table A1.** Incubating parent’s calling rate and fly-off probability *before arrival* of off-duty partner and during *regular* times of incubation.

| Model | Response variable | Effect type | Effect | Estimate | 95% CI |  |
| --- | --- | --- | --- | --- | --- | --- |
|  |  |  |  |  | Lower | Upper |
| a | Calling rate (10min<sup>-1</sup>)<br>(Poisson scale) | Fixed | Intercept (before arrival) | -0.904 | -1.277 | -0.53 |
|  |  |  | **Observation (regular)** | **-0.855** | **-1.12** | **-0.587** |
| b | Calling rate (10min<sup>-1</sup>)<br>(Poisson scale) | Fixed | Intercept (before arrival & ♀) | -0.804 | -1.198 | -0.404 |
|  |  |  | **Observation (regular)** | **-0.88** | **-1.239** | **-0.524** |
|  |  |  | Sex (♂) | -0.18 | -0.453 | 0.091 |
|  |  |  | Day of incubation | -0.083 | -0.307 | 0.152 |
|  |  |  | Sex × Observation | 0.066 | -0.444 | 0.565 |
|  |  |  | Day of incubation × Observation | 0.064 | -0.203 | 0.319 |
| c | Fly-off (0, 1)<br>(binomial scale) | Fixed | Intercept (before arrival) | -2.044 | -2.62 | -1.487 |
|  |  |  | Length of observation (ln(min)) | -0.341 | -0.618 | -0.049 |
|  |  |  | Observation (regular) | -0.195 | -1.03 | 0.651 |
| d | Fly-off (0, 1)<br>(binomial scale) | Fixed | Intercept (before arrival & ♀) | -2.459 | -3.366 | -1.536 |
|  |  |  | Length of observation (ln(min)) | -0.304 | -0.62 | -0.006 |
|  |  |  | Observation (regular) | -0.098 | -1.493 | 1.302 |
|  |  |  | Sex (♂) | 0.739 | -0.429 | 1.882 |
|  |  |  | Day of incubation | 0.015 | -0.497 | 0.527 |
|  |  |  | Sex × Observation | -0.177 | -1.874 | 1.564 |
|  |  |  | Day of incubation × Observation | 0.085 | -0.673 | 0.789 |

Shown are the posterior estimates (medians) of the effect sizes with the 95% credible intervals (CI) from a posterior distribution of 5,000 simulated values generated by the *sim* function in R (Gelman and Su 2018). Variance components were estimated by the *glmer* function in R (Bates *et al.* 2015). *N* = 239 incubation observations (125 of *before arrival*, 114 of *regular*) from 29 nests. Continuous predictors were z-transformed (mean-centred and divided by SD). All four models contained nest ID as random intercept. Calling was entered as count and offset specified as length of observation in minutes (divided by ten minutes and ln-transformed), which not only controlled for the length of observation, but also generated a calling rate 10min<sup>-1</sup>. Distribution of fly-offs was bimodal (22 observations with one fly-off, 2 observations with two fly-offs and 3 with four fly-offs, the remaining observation no fly-offs); thus, in the analyses we used fly-off intensity as binary variable (0 = no fly-off, 1 = at least one fly-off).

**Table A2.**
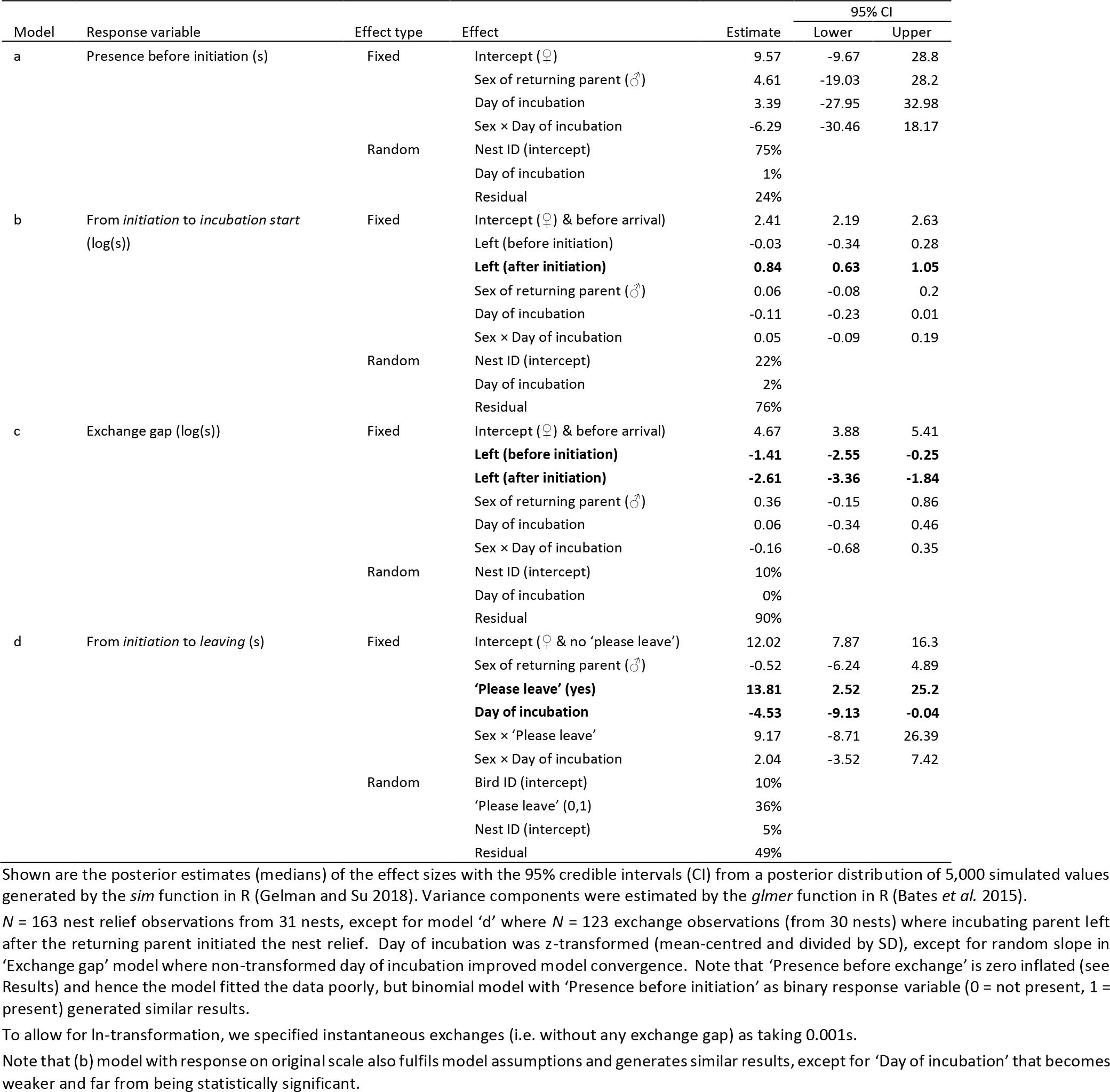
Time components of nest relief behaviour in relation to sex, day of incubation and type of *leaving*.

**Table A3.** ‘Please leave display’ in relation to sex, day of incubation and current bout length.

| Model | Response variable | Effect type | Effect | Estimate | 95% CI |  |
| --- | --- | --- | --- | --- | --- | --- |
|  |  |  |  |  | Lower | Upper |
| a | ‘Please leave’ (0,1)<br>(binomial scale) | Fixed | Intercept (♀) | -3.75 | -5.83 | -1.69 |
|  |  |  | Sex of incubating parent (♂) | 1.31 | -1.28 | 3.78 |
|  |  |  | Current bout (min) | -1.06 | -2.44 | 0.31 |
|  |  |  | Sex × Current bout | 1.57 | -0.26 | 3.39 |
|  |  | Random | Bird ID (intercept) | 99% |  |  |
|  |  |  | Current bout | 1% |  |  |
|  |  |  | Nest ID (intercept) | 0% |  |  |
| b | ‘Please leave’ (0,1)<br>(binomial scale) | Fixed | Intercept (♀) | -6.58 | -11.74 | -1.83 |
|  |  |  | Sex of incubating parent (♂) | 0.49 | -5.84 | 6.88 |
|  |  |  | **Day of incubation** | **1.75** | **0.21** | **3.41** |
|  |  |  | Sex × Day of incubation | 2.59 | -0.95 | 6.01 |
|  |  | Random | Bird ID (intercept) | 93% |  |  |
|  |  |  | Day of incubation | 7% |  |  |
|  |  |  | Nest ID (intercept) | 0% |  |  |

Shown are the posterior estimates (medians) of the effect sizes with the 95% credible intervals (CI) from a posterior distribution of 5,000 simulated values generated by the *sim* function in R (Gelman and Su 2018). Variance components were estimated by the *glmer* function in R (Bates *et al.* 2015). $N_a$ = 117 nest reliefs of 49 birds from 29 nests, $N_b$ = 123 nest reliefs of 51 birds from 30 nests. ‘Current bout’ length and ‘Day of incubation’ were z- transformed (mean-centred and divided by SD).

**Table A4.** Calling during nest relief in relation to sex and day of incubation.

| Model | Response variable | Effect type | Effect | Estimate | 95% CI |  |
| --- | --- | --- | --- | --- | --- | --- |
|  |  |  |  |  | Lower | Upper |
| a | Calling while initiating (0, 1)<br>(binomial scale) | Fixed | Intercept (♀) | -0.885 | -1.48 | -0.31 |
|  |  |  | Sex of returning parent (♂) | -0.287 | -1.161 | 0.595 |
|  |  |  | Day of incubation | 0.015 | -0.663 | 0.686 |
|  |  |  | Sex × Day of incubation | 0.19 | -0.645 | 1.061 |
|  |  | Random | Bird ID (intercept) | 98% |  |  |
|  |  |  | Nest ID (intercept) | 2% |  |  |
| b | Reply of incubating parent (0, 1)<br>(binomial scale) | Fixed | Intercept (♀) | -0.561 | -1.736 | 0.59 |
|  |  |  | Sex of incubating parent (♂) | 0.253 | -1.227 | 1.759 |
|  |  |  | Day of incubation | -0.548 | -1.763 | 0.689 |
|  |  |  | Sex × Day of incubation | 1.105 | -0.637 | 2.844 |
|  |  | Random | Bird ID (intercept) | 100% |  |  |
|  |  |  | Nest ID (intercept) | 0% |  |  |
| c | Calling intensity of returning parent<br>(from initiation to leaving when both<br>parents around) | Fixed | Intercept (♀) | 1.671 | 1.17 | 2.164 |
|  |  |  | Sex of incubating parent (♂) | 0.235 | -0.443 | 0.922 |
|  |  |  | **Calling intensity incubating parent** | **0.496** | **0.112** | **0.894** |
|  |  |  | Day of incubation | 0.164 | -0.253 | 0.578 |
|  |  |  | Sex × Calling incubation parent | -0.523 | -1.052 | 0.011 |
|  |  |  | Sex × Day of incubation | 0.31 | -0.309 | 0.92 |
|  |  | Random | Bird ID (intercept) | 41% |  |  |
|  |  |  | Calling intensity incubating parent | 2% |  |  |
|  |  |  | Nest ID (intercept) | 0% |  |  |
|  |  |  | Residual | 57% |  |  |
| d | Calling intensity during exchange gap | Fixed | Intercept (♀) | 1.685 | 1.316 | 2.039 |
|  |  |  | Sex of returning parent (♂) | -0.378 | -0.873 | 0.143 |
|  |  |  | Day of incubation | 0.219 | -0.173 | 0.61 |
|  |  |  | Sex × Day of incubation | 0.037 | -0.464 | 0.562 |
|  |  | Random | Bird ID (intercept) | 5% |  |  |
|  |  |  | Day of incubation | 3% |  |  |
|  |  |  | Nest ID (intercept) | 0% |  |  |
|  |  |  | Residual | 91% |  |  |
| e | Calling intensity after exchange | Fixed | Intercept (♀) | 0.606 | 0.302 | 0.91 |
|  |  |  | Sex of returned parent (♂) | -0.179 | -0.627 | 0.256 |
|  |  |  | Day of incubation | 0.223 | -0.061 | 0.509 |
|  |  |  | Sex × Day of incubation | -0.131 | -0.533 | 0.253 |
|  |  | Random | Bird ID (intercept) | 43% |  |  |
|  |  |  | Calling during return | 8% |  |  |
|  |  |  | Nest ID (intercept) | 0% |  |  |
|  |  |  | Residual | 49% |  |  |

Shown are the posterior estimates (medians) of the effect sizes with the 95% credible intervals (CI) from a posterior distribution of 5,000 simulated values generated by the *sim* function in R (Gelman and Su 2018). Variance components were estimated by the *lmer* or *glmer* functions in R (Bates *et al.* 2015).
Predictors (Day of incubation and Calling intensity) were z-transformed (mean-centred and divided by SD). Note that effects of those continuous predictors that are not also specified as random slopes (because of convergence issues) might be inflated.
$N_{\text{Calling while initiating}}$ = 117 nest reliefs (with incubating parent leaving only after partner initiated the nest relief) of 47 parents from 28 nests.
$N_{\text{Reply of incubating parent}}$ = 31 nest reliefs (with returning parent calling while arriving) of 24 parents from 19 nests.
$N_{\text{Calling intensity of returning parent}}$ = 89 nest reliefs of 42 parents from 26 nests.
$N_{\text{Calling intensity during exchange gap}}$ = 118 nest reliefs of 48 parents from 29 nests.
$N_{\text{Calling intensity after exchange}}$ = 117 nest reliefs of 47 parents from 28 nests.

**Table A5.** Probability of calling while leaving the nest.

| Response variable | Effect type | Effect | Estimate | 95% CI |  |
| --- | --- | --- | --- | --- | --- |
|  |  |  |  | Lower | Upper |
| Called when left (0,1)<br>(binomial scale) | Fixed | Intercept (♀) | -2.22 | -3.04 | -1.41 |
|  |  | **Sex of incubating parent (♂)** | **1.01** | **0.08** | **1.95** |
|  |  | Day of incubation (min) | -0.3 | -1.13 | 0.54 |
|  |  | Sex × Day of incubation | 0.24 | -0.76 | 1.21 |
|  | Random | Bird ID (intercept) | 13% |  |  |
|  |  | Day of incubation | 22% |  |  |
|  |  | Nest ID (intercept) | 66% |  |  |

Shown are the posterior estimates (medians) of the effect sizes with the 95% credible intervals (CI) from a posterior distribution of 5,000 simulated values generated by the *sim* function in R (Gelman and Su 2018). Variance components were estimated by the *glmer* function in R (Bates *et al.* 2015). *N* = 150 nest reliefs of 55 birds from 30 nests. ‘Day of incubation’ was z-transformed (mean-centred and divided by SD). A model without the interaction generated similar results.

**Table A6.**
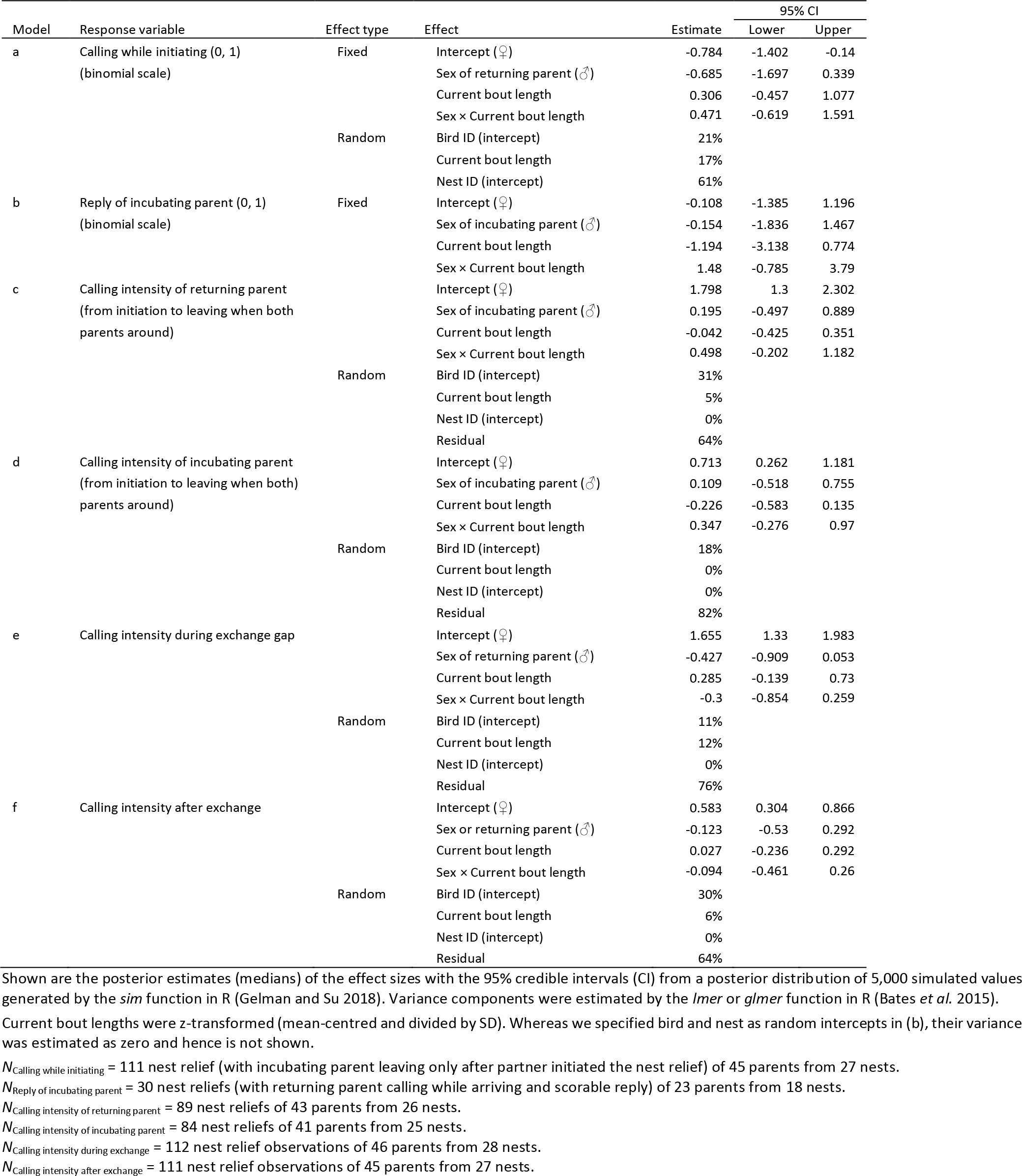
Calling during nest relief in relation to current bout length.

**Table A7.** Next incubation bout in relation to calling during nest relief.

| Model | Response variable | Effect type | Effect | Estimate | 95% CI |  |
| --- | --- | --- | --- | --- | --- | --- |
|  |  |  |  |  | Lower | Upper |
| a | Next bout (min) | Fixed | Intercept (♀ & no arrival call) | 685 | 627.5 | 743.9 |
|  |  |  | Arrival call (yes) | 44.1 | -36.4 | 123 |
|  |  |  | Sex of returning parent (♂) | -25.5 | -104.8 | 52.2 |
|  |  |  | Sex × Arrival call | -16.4 | -130 | 95.3 |
|  |  | Random | Bird ID (intercept) | 39% |  |  |
|  |  |  | Nest ID (intercept) | 4% |  |  |
|  |  |  | Residual | 57% |  |  |
| b | Next bout (min) | Fixed | Intercept (♀ & no reply call) | 596.1 | 504.3 | 690.3 |
|  |  |  | **Reply call (yes)** | **251.7** | **93.9** | **405.6** |
|  |  |  | Sex of incubating parent (♂) | 85.2 | -44.6 | 210.3 |
|  |  |  | Sex × Reply call | -158.9 | -364.4 | 54.8 |
|  |  |  | Bird ID (intercept) | 19% |  |  |
|  |  |  | Nest ID (intercept) | 4% |  |  |
|  |  |  | Residual | 77% |  |  |
| c | Next bout (min) | Fixed | Intercept (♀) | 698.8 | 636.1 | 759.5 |
|  |  |  | Sex of returning parent (♂) | -20.7 | -101.6 | 59.3 |
|  |  |  | Calling intensity returning parent | 16.3 | -35.3 | 67.3 |
|  |  |  | Calling intensity incubating parent | 19.6 | -24.8 | 62.5 |
|  |  |  | Sex × Calling intensity returning parent | -16.8 | -87.7 | 53.4 |
|  |  |  | Sex × Calling intensity incubating parent | 11.6 | -54.9 | 77.4 |
|  |  | Random | Bird ID (intercept) | 22% |  |  |
|  |  |  | Nest ID (intercept) | 13% |  |  |
|  |  |  | Residual | 65% |  |  |
| d | Next bout (min) | Fixed | Intercept (♀ & no left call) | 686.6 | 624.7 | 746.3 |
|  |  |  | Sex of incubating parent (♂) | -13.9 | -96.2 | 72 |
|  |  |  | Called when left (yes) | -41.6 | -138.2 | 55.1 |
|  |  |  | Sex × Called when left | 9.7 | -126 | 151 |
|  |  | Random | Bird ID (intercept) | 47% |  |  |
|  |  |  | Nest ID (intercept) | 0% |  |  |
|  |  |  | Residual | 53% |  |  |
| e | Next bout (min) | Fixed | Intercept (♀) | 676.7 | 615.6 | 732.8 |
|  |  |  | Sex of returning parent (♂) | -2.7 | -82.4 | 81 |
|  |  |  | Calling during exchange gap | 21 | -14.2 | 55.4 |
|  |  |  | Sex × Calling during exchange gap | 14.3 | -36 | 64.3 |
|  |  | Random | Bird ID (intercept) | 50% |  |  |
|  |  |  | Nest ID (intercept) | 2% |  |  |
|  |  |  | Residual | 48% |  |  |
| f | Next bout (min) | Fixed | Intercept (♀) | 668.5 | 604.8 | 731 |
|  |  |  | Sex of returning parent (♂) | -7.5 | -83.1 | 69.2 |
|  |  |  | **Calling after incubation start** | **49.9** | **6.7** | **91.6** |
|  |  |  | **Sex × Calling after incubation start** | **-66.6** | **-127** | **-0.9** |
|  |  | Random | Bird ID (intercept) | 38% |  |  |
|  |  |  | Calling after incubation start | 11% |  |  |
|  |  |  | Nest ID (intercept) | 20% |  |  |
|  |  |  | Residual | 31% |  |  |

Shown are the posterior estimates (medians) of the effect sizes with the 95% credible intervals (CI) from a posterior distribution of 5,000 simulated values generated by the *sim* function in R (Gelman and Su 2018). Variance components were estimated by the *lmer* function in R (Bates *et al.* 2015).
Calling intensities were z-transformed (mean-centred and divided by SD). Because in (f) calling seemed statistically supported, to account for possible non-independence of data points, we have included calling also as a random slope (Schielzeth and Forstmeier 2009). Included are only nest relief where incubating parent left after its partner initiated the nest relief.
$N_a$ = 104 nest reliefs of 44 parents from 26 nests.
$N_b$ = 28 nest reliefs (with returning parent calling while arriving and scorable reply) of 22 parents from 17 nests.
$N_c$ = 79 nest reliefs of 39 parents from 24 nests.
$N_d$ = 101 nest reliefs of 45 parents from 25 nests.
$N_e$ = 105 nest reliefs of 45 parents from 27 nests.
$N_f$ = 104 nest reliefs of 44 parents from 26 nests.

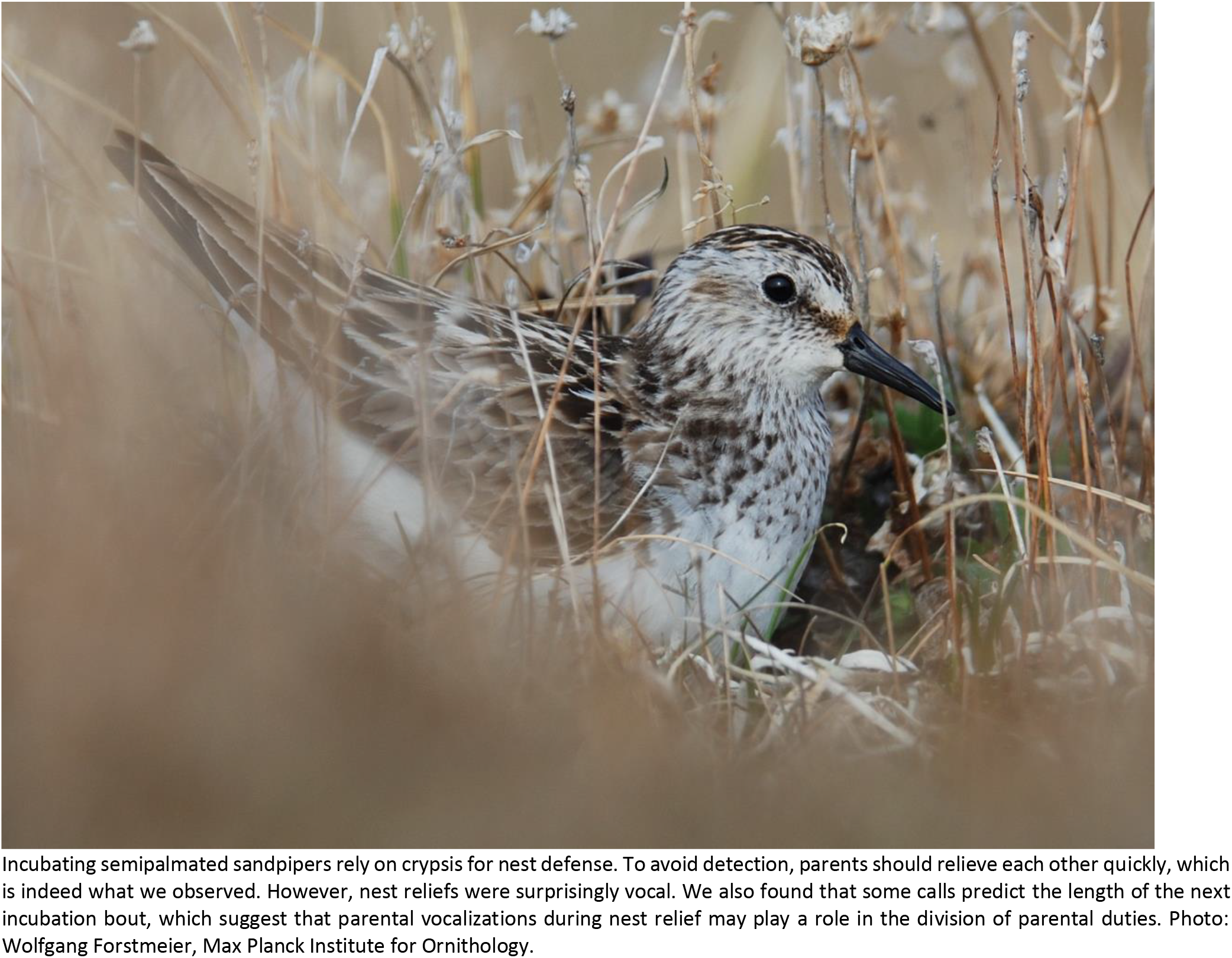
GRAPHICAL ABSTRACT.

